# RSim: A Reference-Based Normalization Method via Rank Similarity

**DOI:** 10.1101/2023.04.04.535620

**Authors:** Bo Yuan, Shulei Wang

## Abstract

Microbiome sequencing data normalization is crucial for eliminating technical bias and ensuring accurate downstream analysis. However, this process can be challenging due to the high frequency of zero counts in microbiome data. We propose a novel reference-based normalization method called normalization via rank similarity (RSim) that corrects sample-specific biases, even in the presence of many zero counts. Unlike other normalization methods, RSim does not require additional assumptions or treatments for the high prevalence of zero counts. This makes it robust and minimizes potential bias resulting from procedures that address zero counts, such as pseudo-counts. Our numerical experiments demonstrate that RSim reduces false discoveries, improves detection power, and reveals true biological signals in downstream tasks such as PCoA plotting, association analysis, and differential abundance analysis.

## 1 Introduction

High-throughput sequencing technology has revolutionized the study of microbiome communities, providing biologists with a powerful tool to investigate and understand biological events and mechanisms. However, analyzing and interpreting the data generated by high-throughput sequencing can be challenging due to technical factors that can confound results (Vallejos et al., 2017; Weiss et al., 2017; Lin and Peddada, 2020). One major limitation of high-throughput sequencing is that the observed sequencing count data can only reflect the relative abundance of taxa rather than their absolute abundance, as the observed sequencing depth can vary significantly across samples and is unrelated to the absolute abundance (Robinson and Oshlack, 2010; Young et al., 2010; Conesa et al., 2016). In mathematical terms, the observed sequencing count data can be expressed as:

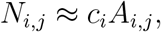

where *N*_*i,j*_ and *A*_*i,j*_ are the observed count and absolute abundance of taxon *j* in the *i*th sample, and *c*_*i*_ is the unobserved sampling fraction of the *i*th sample. The unobserved sampling fraction is typically sample-specific and can vary due to technical factors such as sequencing depth and capture efficiency. Because of this unobserved sampling fraction, applying classical statistical methods to the observed count data can result in false-positive scientific discoveries and invalid analysis results (Vandeputte et al., 2017; Weiss et al., 2017). This paper refers to the bias resulting from the unobserved sampling fraction as compositional bias.

Normalizing the observed sequencing count data is a critical step in removing compositional bias and ensuring accurate and reliable downstream analysis. To this end, many normalization methods have been proposed for different types of sequencing data sets (Hughes and Hellmann, 2005; Robinson and Oshlack, 2010; Anders and Huber, 2010; Bullard et al., 2010; Dillies et al., 2013; Paulson et al., 2013; Lun et al., 2016; Bacher et al., 2017; Kumar et al., 2018; Hafemeister and Satija, 2019). These methods can be broadly classified into three computational frameworks: rarefying, scaling, and log-ratio based methods (Weiss et al., 2017; Lin and Peddada, 2020). The rarefying method subsamples the taxa of each sample to ensure that all samples have the same sequencing depth. Although this method is popular in practice, it may lead to a loss of statistical power in downstream analysis and does not correct compositional bias (McMurdie and Holmes, 2014). Besides rarefying method, the scaling method is another widely used normalization strategy that estimates the unobserved sampling fraction and scales the observed count by this estimated sampling fraction. Scaling methods include Cumulative-Sum Scaling (CSS) (Paulson et al., 2013), Median (MED) (Love et al., 2014), Upper Quartile (UQ) (Bullard et al., 2010), Trimmed Mean of M-values (TMM) (Robinson and Oshlack, 2010), and Total-Sum Scaling (TSS) normalization. However, accurately estimating the sampling fraction can be challenging when prevalent zero counts exist in the microbiome data (Weiss et al., 2017). Finally, log-ratio based methods, which are motivated by classical compositional data analysis (Aitchison, 1982; Pawlowsky-Glahn and Buccianti, 2011), have been proposed. Although log-ratio transformation can alleviate the compositional effect, it is still unclear how to apply log-ratio transformation when zero counts are present and how to interpret the results (Greenacre, 2021; Brill et al., 2022; Wang, 2023b). These challenges lead us to question whether a new normalization method can be developed that is both robust to the prevalent zero counts and corrects the compositional bias.

Here, we introduce a novel normalization method, which we call RSim (normalization via *R*ank *Sim*ilarity), to correct the compositional bias in the sequencing data set. The RSim normalization is a scaling method motivated by the normalization method in the experiment with spike-in bacteria. Instead of estimating sampling fraction directly, RSim first identifies a set of non-differential abundant taxa via the pairwise rank similarity of taxa and then scales the counts to ensure that the total sum of coverage in this estimated set is the same across samples. To accurately identify non-differential abundant taxa, RSim employs a new empirical Bayes approach to control the misclassification rate. Unlike existing methods, RSim does not need any assumption or extra treatment to the prevalent zero counts because the Spearman ‘s rank correlation coefficient used for measuring rank similarity is robust to zero counts. Besides being robust to zero, RSim outperforms existing methods in estimating the sampling fraction and correcting the compositional bias. We demonstrate the efficacy of RSim by comparing it with several state-of-the-art methods using synthetic and real data sets. Our results show that RSim can help reduce false discoveries, improve detection power, and reveal true biological signals in various downstream analyses, such as PCoA plotting, association analysis, and differential abundance analysis. RSim normalization is implemented in an R package, freely available at https://github.com/BoYuan07/RSimNorm.

## 2 Results

### 2.1 Overview of RSim Normalization

We present a concise summary of the RSim normalization method, with a more detailed explanation provided in the Method section. While we focus on the microbiome data in this paper, it is worth noting that RSim may be applicable to other sequencing data, such as bulk RNA-seq and single-cell RNA-seq (Vallejos et al., 2017). The RSim method is inspired by the normalization approach used in experiments with spike-in bacteria (Stämmler et al., 2016; Tourlousse et al., 2017; Tkacz et al., 2018; Hardwick et al., 2018). When spike-in bacteria are available, the count of each taxon is rescaled by the reciprocal of the count of the spike-in taxa, as follows:

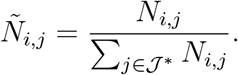

Here, *N*_*i,j*_ represents the observed count of taxon *j* in the *i*th sample, and 𝒥* is the set of spike-in taxa. We refer to this method as reference-based normalization since it treats 𝒥* as a reference set. The reference-based method can correct compositional bias when spike-in bacteria are available (Stämmler et al., 2016). The efficacy of the reference-based method is contingent on the assumption that the absolute abundance of spike-in taxa is identical across samples, as expressed by the equation:

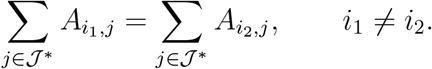

Here, *A*_*i,j*_ represents the absolute abundance of taxon *j* in the *i*th sample. Given the success of the spike-in based normalization, one might wonder whether it is feasible to identify a reference set of taxa that satisfies the above equality and use it to correct compositional bias without employing spike-in bacteria. Our paper shows that this is possible when we can identify a set of non-differential abundant taxa, denoted by 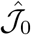, and replace the spike-in taxa with these estimated non-differential abundant taxa.

The RSim normalization method is primarily aimed at identifying a set of non-differential abundant taxa in microbiome data, even in the presence of zero counts. This identification process has two steps: first, we construct statistics for the differential abundance level of each taxon by using pairwise rank similarity of taxa; second, we use a new empirical Bayes method to identify non-differential abundant taxa based on the statistics obtained in the first step (see Figure 1). To explain the intuition behind the first step, we note that the count of a non-differential abundant taxon is approximately proportional to the unknown sampling fraction, whereas the count of a differential abundant taxon lacks this correspondence to the sampling fraction. Thus, the rank correlation between two non-differential abundant taxa should be higher than between a non-differential abundant taxon and a differential abundant taxon. Assuming that the majority of taxa are non-differential abundant, we use the median of rank correlation coefficients between a taxon and other taxa as the statistics for the level of differential abundance. In the second step, we use an empirical Bayes method to identify non-differential abundant taxa based on the statistics obtained in the first step. The new empirical Bayes method allows choosing a threshold to control the misclassification error. Since most identified taxa are non-differential abundant, they can serve as the reference set for the reference-based normalization in RSim. Notably, the RSim procedure treats zero entries the same way as non-zero entries, allowing it to work consistently with zero counts. In the following sections, we demonstrate the effectiveness of RSim normalization in correcting compositional bias, even in the presence of many zeros in the data set.

**Figure 1:**
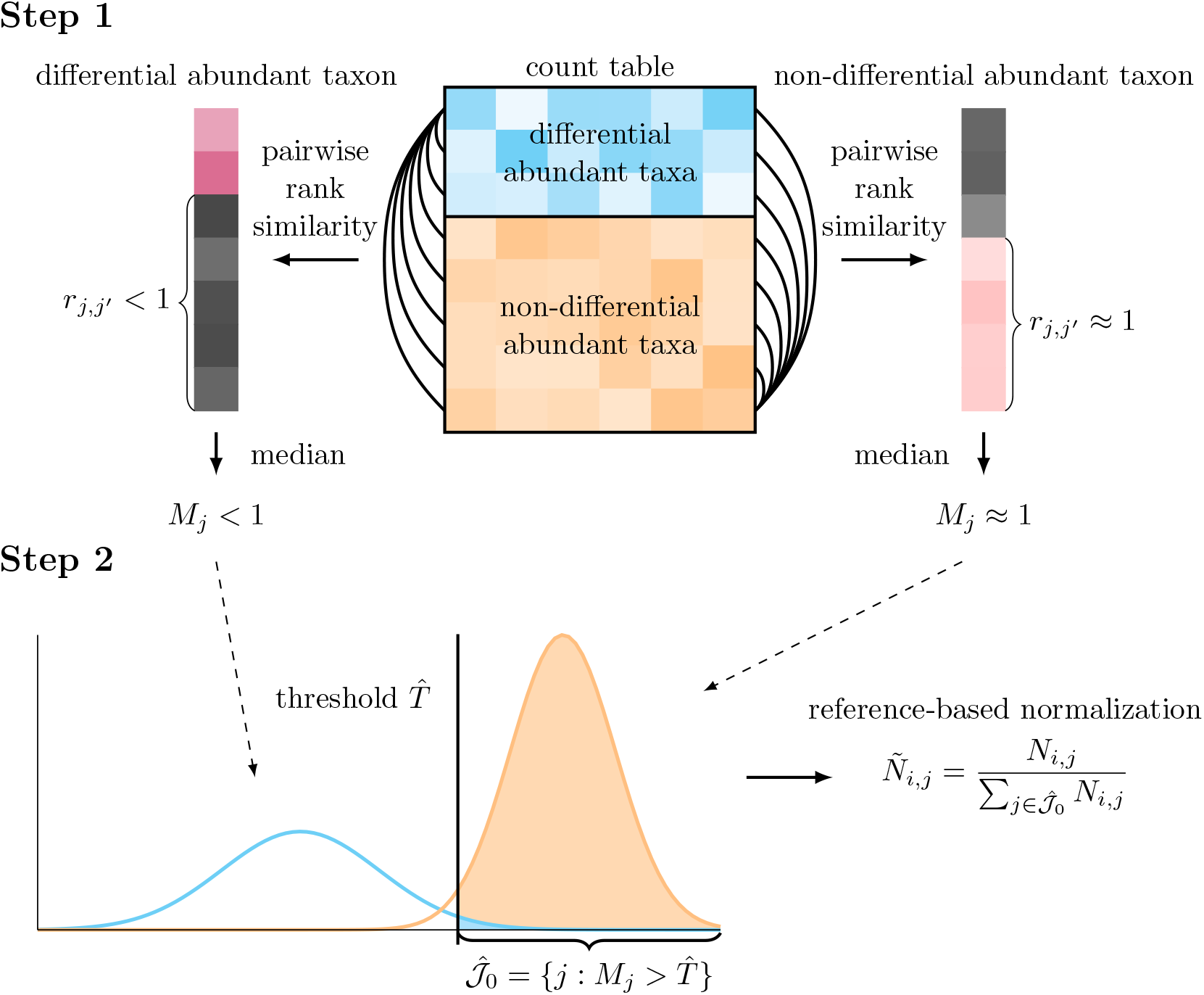
Illustration demonstrating the procedure of RSim normalization. Step 1: median of pairwise rank similarity of taxa is evaluated to construct the statistics for the differential abundance level of each taxon. Step 2: a new empirical Bayes method provides misclassification rate control in identifying non-differential abundant taxa. Estimated non-differential abundant taxa are used as the reference set in reference-based normalization.

### 2.2 Correcting Compositional Bias via RSim Normalization

This section presents a series of numerical experiments to assess the ability of RSim normalization to correct compositional bias. We generate synthetic data using a microbiome dataset collected in He et al. (2018), where 97% of entries are zeros. We first investigate whether the taxa in the estimated reference set of RSim normalization are mostly non-differentially abundant. Specifically, we design several numerical experiments to determine if the empirical Bayes method in RSim can control the misclassification rate in the estimated reference set at a desired level. Figure S1 shows that RSim successfully controls the empirical misclassification rate at the target level for different levels of misclassification rate. We further evaluate the robustness of the estimated reference set by varying the signal strength of differential abundant taxa, the balance of group size in differential abundant taxa, the proportion of differential abundant taxa, and sample size change (Figure S2). Our experiments demonstrate that RSim can robustly identify a reference set that consists mostly of non-differential abundant taxa.

The next set of numerical experiments aims to investigate whether RSim normalization can recover the sampling fraction of each sample via reference-based normalization. To this end, we compare RSim normalization with five state-of-art normalization methods, including TSS, UQ implemented in edgeR, CSS implemented in metagenomeSeq, MED implemented in DESeq2, and TMM implemented in edgeR. We also include an oracle reference-based normalization where the reference set consists of true non-differential abundant taxa. Using synthetic data generated from a microbiome dataset collected in He et al. (2018), we randomly divided the samples into two groups and inserted signals to the differential abundant taxa of one group. We then estimated the sampling fraction of each sample using the seven normalization methods and compared their performance. Figure 2 presents the results of these experiments. We found that when the signal strength of differential abundant taxa is weak, most normalization methods can recover the sampling fraction well and do not exhibit significant bias in their estimates. However, in the presence of strong differential abundant taxa, existing normalization methods suffer from a systematic bias in sampling fraction estimation, while reference-based normalization is robust to this bias. Notably, RSim normalization performs similarly to the oracle method, indicating that the reference set selected by RSim contains mostly non-differential abundant taxa and can effectively normalize the data. Overall, our numerical experiments demonstrate that RSim normalization corrects the sample-specific bias resulting from technical variations in the sequencing process. In the next three sections, we investigate how RSim normalization improves the performance of commonly used downstream analyses, including PCoA plotting, association analysis, and differential abundance analysis.

**Figure 2:**
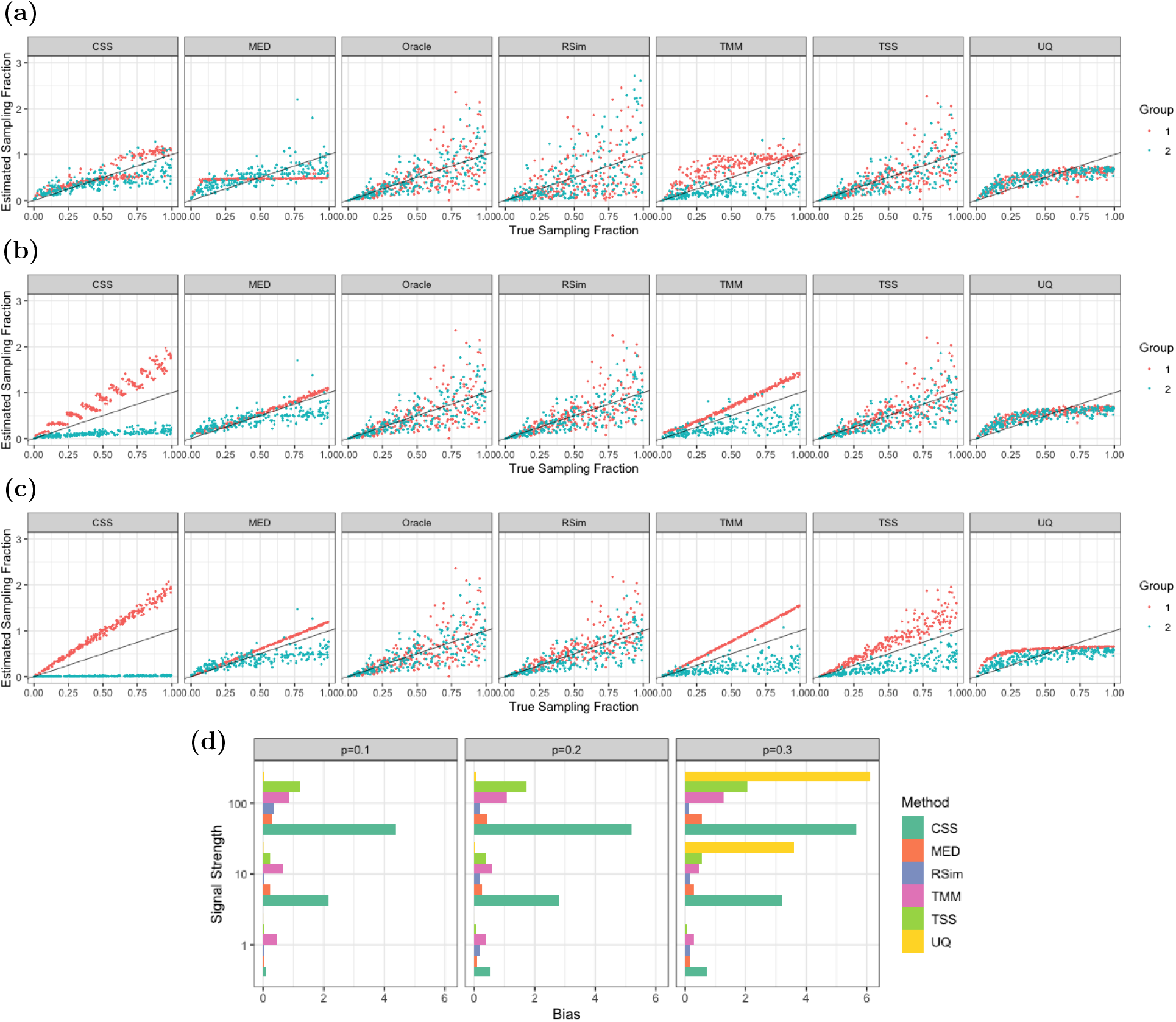
Comparisons of normalization methods in estimating sampling fraction. The numerical experiments are performed when the signal strength of differential abundant taxa is (a) weak, (b) moderate, and (c) strong. In (a), (b), and (c), the *x*-axis represents true sampling fractions, while the *y*-axis represents the estimated sampling fraction from normalization methods. We scale the estimated sampling fractions so that their average is the same as the average of true sampling fractions. The black line in these figures represents equality between the estimated and true sampling fractions and the color of points represent which group the differential abundant taxa belong to. The bias in sampling fraction estimation by different normalization methods is compared in (d) when the signal strength and proportion (*p* = 0.1, 0.2, 0.3) of differential abundant taxa vary. It is clear that the reference-based method can better correct the compositional bias than existing methods, especially when there is a large proportion of strong differential abundant taxa.

### 2.3 RSim Normalization Reveals Biological Pattern in PCoA Plot

This section aims to investigate the effects of different normalization methods on the PCoA plots. Specifically, we compare the PCoA plots on the normalized data after applying six normalization methods, namely TSS, UQ, CSS, MED, TMM, and RSim, to a microbiome data set collected in He et al. (2018). The compositional bias can create false clusters or patterns in the PCoA plot if the count data is not appropriately normalized. We randomly split the samples into two groups, and no cluster structure is observed in the PCoA plots, regardless of the normalization method used (Figure 3a). However, when we rarefy the count data of one group of samples through subsampling, the difference in the sequencing depth results in two clusters in some of the PCoA plots (Figure 3b). In particular, RSim, TMM, and TSS can remove such false clusters through normalization, while CSS, MED, and UQ cannot. We also conduct a similar numerical experiment on another dataset collected in Vangay et al. (2018). The samples in the KarenThai category are divided into two groups based on the sequencing depth (>10000 belongs to the first group, and <5000 belongs to the second group). The two clusters separated by sequencing depth are present in the PCoA plots of all normalization methods except RSim normalization (Figure 3c). Through these two examples, we conclude that RSim normalization is more effective in mitigating the issue of false clusters or patterns in PCoA plots than existing normalization methods.

**Figure 3:**
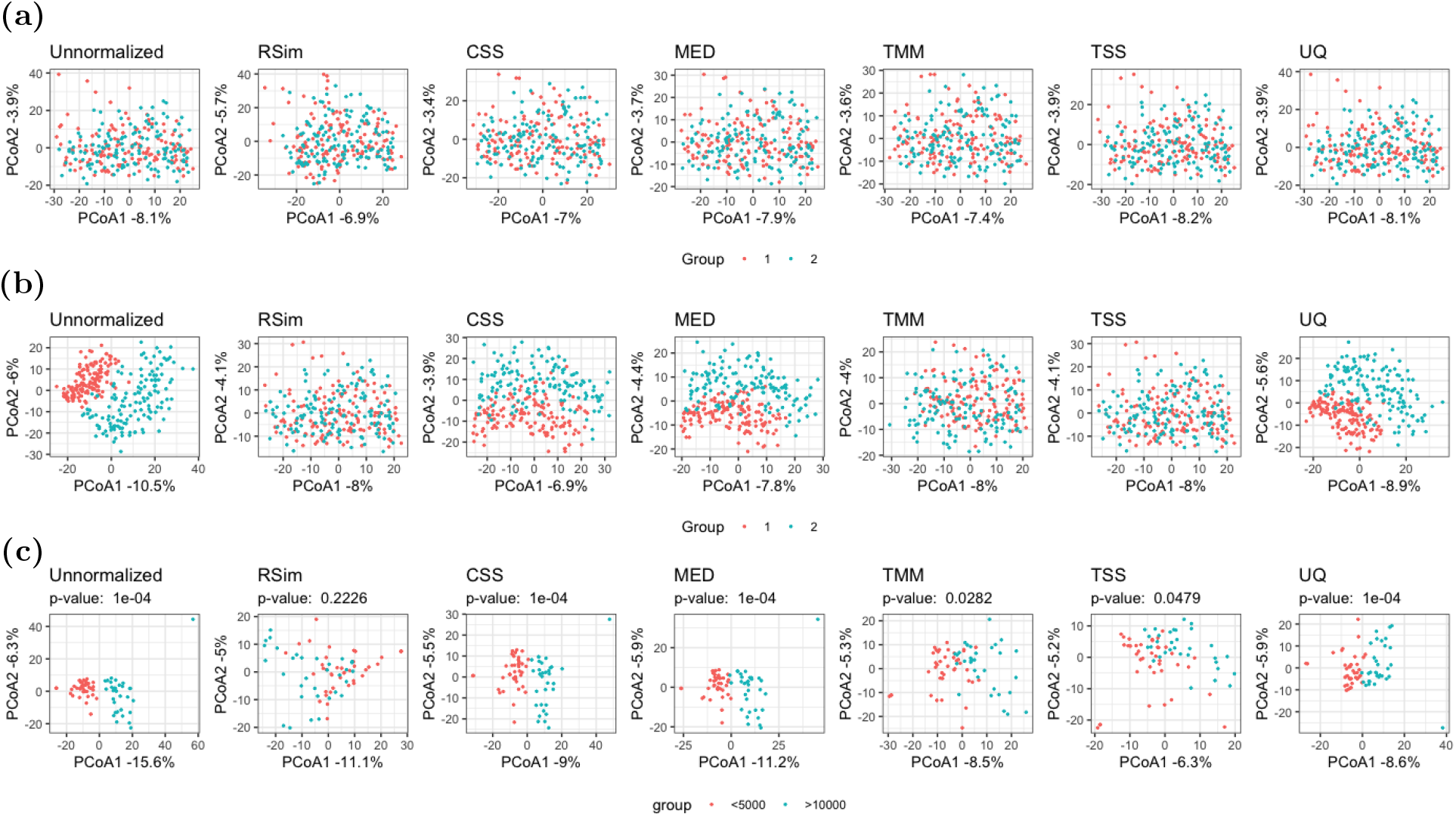
Compositional bias can create false clusters in PCoA plots. In (a) and (b), samples are randomly divided into two groups. No modification is applied to (a), while the count data in group 1 is rarefied in (b). In (c), samples are divided into two groups based on the sequencing depth (>10000 belongs to the first group, and <5000 belongs to the second group). In these figures, RSim normalization can help remove the false clusters resulting from compositional bias. Euclidean distance with log transformation is used in all PCoA plots.

The presence of false patterns resulting from compositional bias can lead to erroneous interpretations of data, highlighting the importance of proper normalization. Figure S4a shows the PCoA plots of right palm samples in a data set collected in Caporaso et al. (2011), colored by days since the experiment started. The PCoA plot exhibits a clear time-related pattern in the raw data, implying a possible shift in microbial abundance during the 15-month study period. Similar patterns are also observed in the PCoA plots after applying all normalization methods except RSim. However, further examination reveals a strong correlation between time and sequencing depth (Figure S4c), and a similar pattern is also present in the PCoA plots colored by sequencing depth (Figure S4b). This suggests that sequencing depth is a confounding factor and is likely responsible for the observed time-related pattern. RSim normalization can effectively remove this false pattern, and as a result, the pattern of time and sequencing depth in the PCoA plots is no longer apparent, demonstrating the effectiveness of RSim normalization in removing confounding effects.

Appropriate normalization not only helps avoid false clusters but also aids in detecting true biological patterns resulting from microbial abundance shifts. Following a similar approach as in Section 2.2, we generated data with differentially abundant taxa from a data set in He et al. (2018). The absolute abundance of these taxa depends on a latent variable characterizing the biological structure. When this latent variable is binary, a two-cluster structure is expected in the PCoA plot, but compositional bias confounds such a structure (Figure S3a). After applying normalization methods, only RSim normalization helps detect a two-cluster structure in the PCoA plot. We observed a similar phenomenon in the numerical experiment when the latent variable is continuous (Figure S3b). These examples suggest that compositional bias can obscure biological signals of interest, and RSim normalization can help reveal the true biological pattern in the data set.

### 2.4 RSim Normalization Increases Efficiency of Association Analysis

This section investigates the impact of normalization on association analysis, which aims to detect an association between microbiome data and a specific outcome, such as age or BMI. To compare the performance of different normalization methods, we consider two commonly used association analysis methods, PERMANOVA (McArdle and Anderson, 2001; Wang et al., 2021) and MiRKAT (Zhao et al., 2015). Similar to the previous sections, we generate synthetic data from the microbiome data set in He et al. (2018). In the first set of experiments, we examine the effect of normalization on the type I error of association analysis. We randomly divide samples into two groups and rarefy the first group via subsampling. The type I error is highly inflated when we directly apply association analysis to the unnormalized count data due to the difference in sequencing depth. After applying six different normalization methods, only TSS and RSim normalization can effectively control the type I error. We also apply PERMANOVA to samples of Karen individuals living in Thailand, as collected in Vangay et al. (2018), using the same experiment settings as in the previous section. The *P*-values are reported in Figure 3c. PERMANOVA finds a significant association between the microbiome data and the group defined by the sequencing depth when the count data is normalized by the existing normalization method. However, the association is no longer significant when we apply RSim normalization, indicating that RSim normalization can correct the compositional bias resulting from the confounding sequencing depth. These results further confirm that RSim normalization can reduce false discoveries in association analysis.

The second set of numerical experiments investigates the effect of different normalization methods on the power of association analysis. As in the previous two sections, we generated synthetic data with differential abundant taxa from the microbiome data set in He et al. (2018). Applying PERMANOVA and MiRKAT directly to the unnormalized data resulted in power loss, while RSim normalization improved their power more effectively than existing methods in most settings (see Figures 4b and S5). In addition to the synthetic data, we also compared different normalization methods using the data set collected in Vangay et al. (2018). Specifically, we applied PERMANOVA and MiRKAT to examine the global association between BMI and the human gut microbiome in Karen individuals living in Thailand. When the microbiome data was normalized using RSim, we observed a significant association with *P*-values smaller than 0.05. However, when other existing normalization methods were used, no significant discovery was reported (see Table 1). This discovery aligns with previous literature that shows the significant impact of gut microbiota on nutrient metabolism and energy expenditure (Aoun et al., 2020). These findings highlight the importance of appropriate normalization in association analysis to avoid false discoveries and improve power, and RSim normalization is a superior choice to existing methods.

**Figure 4:**
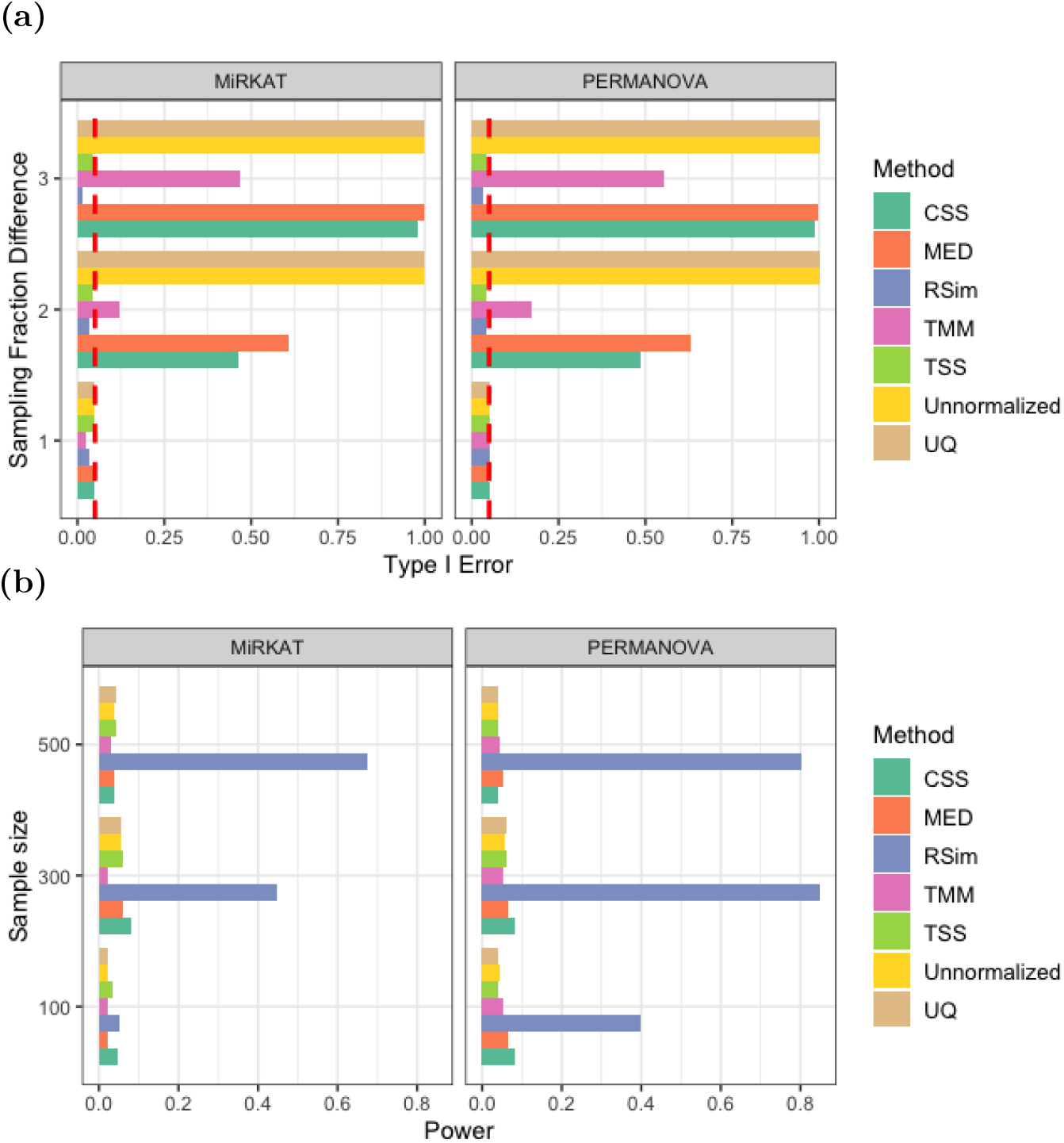
Normalization can reduce false discovery and improve the power of association analysis. In (a), the samples are randomly divided into two groups, and the count data in the first group is rarefied. In (b), the synthetic data include differential abundant taxa. The significance level is 0.05 in both and (b). Normalization is an essential step to avoid false discovery and improve power.

**Table 1:**
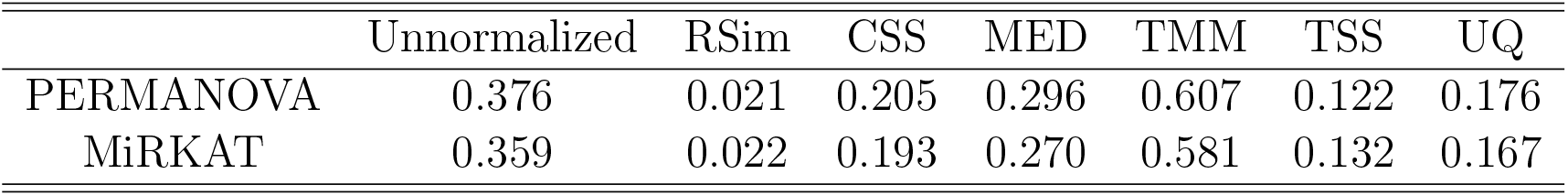
Normalization can make more scientific discoveries through improving the power of association analysis. *P*-values of PERMANOVA and MiRKAT are reported in the study of association between the gut microbiome and BMI.

|  | Unnormalized | RSim | CSS | MED | TMM | TSS | UQ |
| --- | --- | --- | --- | --- | --- | --- | --- |
| PERMANOVA | 0.376 | 0.021 | 0.205 | 0.296 | 0.607 | 0.122 | 0.176 |
| MiRKAT | 0.359 | 0.022 | 0.193 | 0.270 | 0.581 | 0.132 | 0.167 |

### 2.5 RSim Normalization Improves Accuracy of Differential Abundance Analysis

This section focuses on the effect of normalization on differential abundance analysis, which aims to identify taxa with different abundances across conditions. Classical tests, such as the two-sample *t*-test and Pearson correlation test, are commonly used for this analysis, but applying them directly to unnormalized count data can lead to inflated false discoveries. Proper normalization is, therefore, essential to mitigate this issue. We conducted experiments on synthetic data generated from the dataset in He et al. (2018) to study how the normalization impacts differential abundance analysis. Specifically, we apply six normalization methods (TSS, UQ, CSS, MED, TMM, and RSim) and conduct differential abundance analysis using the two-sample *t*-test for binary outcomes and Pearson correlation test for continuous outcomes. The results in Figure 5 showed that inappropriate normalization could introduce bias, resulting in an inflated false discovery rate (FDR) and reduced power. However, RSim normalization was effective in controlling FDR and maintaining sufficient power, thereby mitigating compositional bias. These results confirm the importance of appropriate normalization for differential abundance analysis and suggest that RSim normalization is a reliable choice.

**Figure 5:**
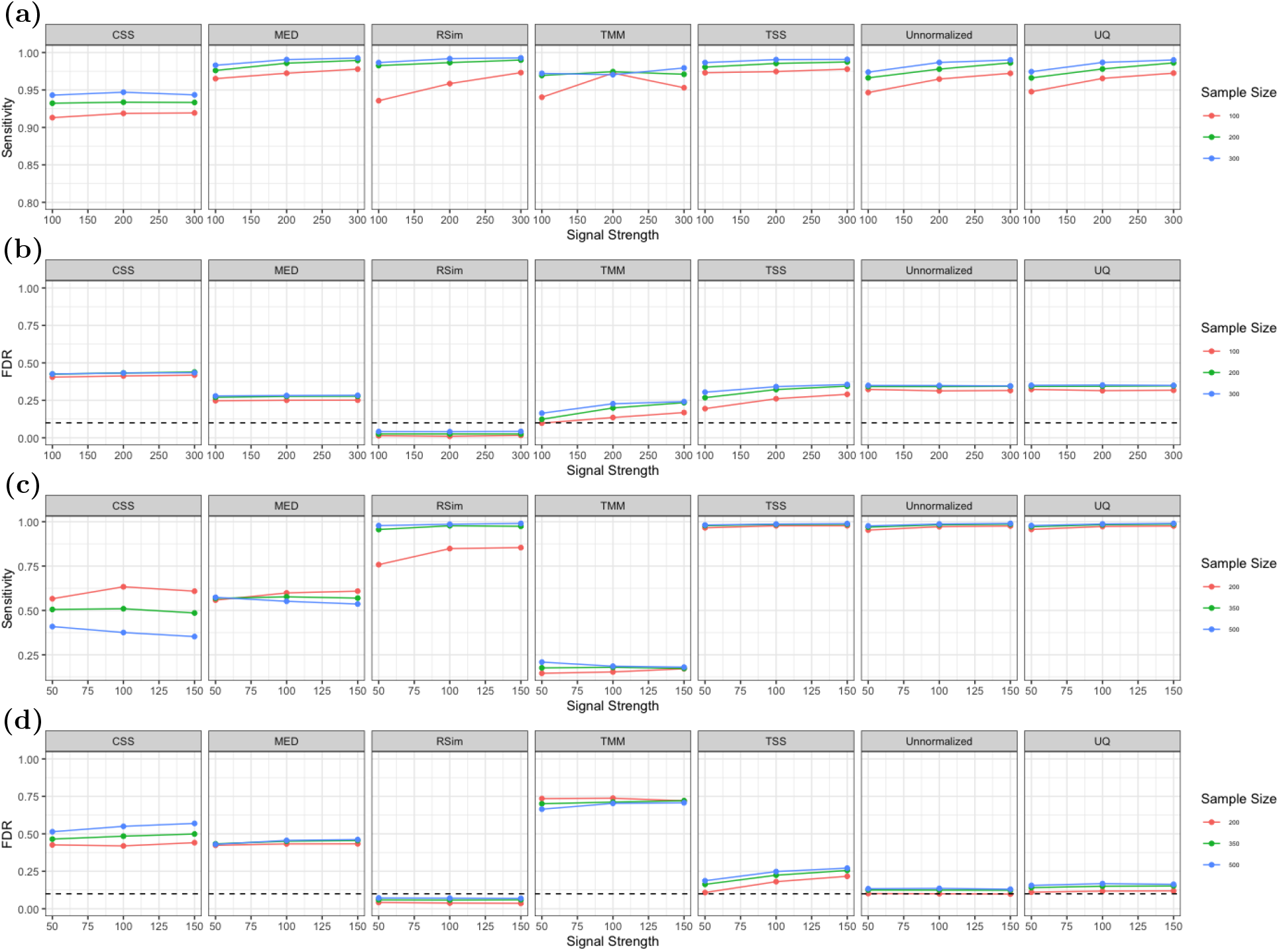
Comparison of different normalization methods’ effect on the differential abundance analysis. (a) and (b) are the FDR and sensitivity plots of the *t*-test after applying six normalization methods. (c) and (d) are the FDR and sensitivity plots of the Pearson correlation test after applying six normalization methods. The *x*-axis is the signal strength of differential abundant taxa. RSim can help *t*-test and Pearson correlation test control FDR and maintain detection power.

In addition to the synthetic data, we also applied RSim normalization to real datasets to further elucidate the effect of normalization on differential abundance analysis. First, we used the dataset from Caporaso et al. (2011) to compare the six normalization methods. The samples were divided into two groups based on sequencing depth, and we applied the two-sample *t*-test equipped with six normalization methods as well as four state-of-the-art differential abundance tests designed for compositional data: ANCOM (Mandal et al., 2015), edgeR (Robinson et al., 2010), LinDA (Zhou et al., 2022), and RDB (Wang, 2023b). The results, as summarized in Figure 6, showed that inappropriate normalization could lead to inflated FDR when the sequencing count data is analyzed. However, RSim normalization successfully corrected for compositional bias and improved the two-sample *t*-test to control FDR and detect significant differences.

**Figure 6:**
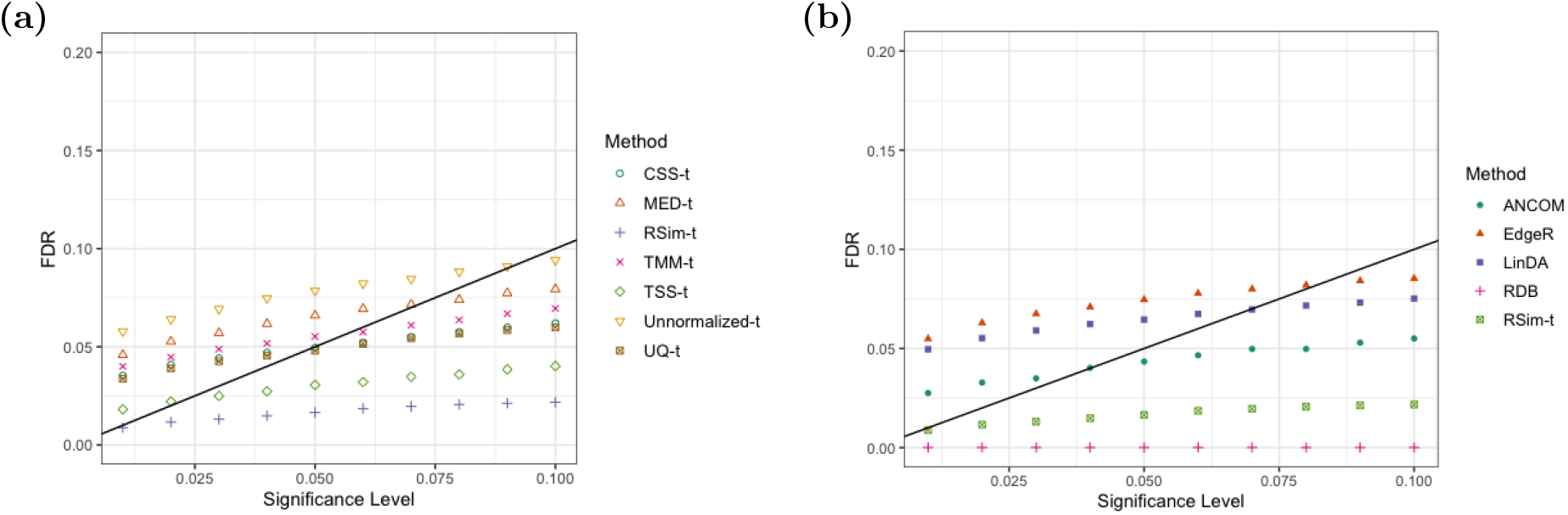
RSim normalization helps two-sample *t*-test control false discovery. Samples are divided into two groups based on the sequencing depth (<10000 belongs to the first group, and >20000 belongs to the second group), and the FDR is shown when the different significance levels are used. In (a), six normalization methods are compared. In (b), a two-sample *t*-test equipped with RSim normalization is compared with state-of-art differential abundance tests.

We also applied RSim normalization and the two-sample *t*-test to a gut microbiome data set from an immigration effect study (Vangay et al., 2018). This analysis compared two groups: Karen (Karen female individuals living in Thailand) vs. Karen1st (Karen female individuals born in Southeast Asia and moved to the US). We detected six significant phyla in the comparison (Table S1). It is noteworthy that the RDB test, which is designed to remove compositional bias and has the best false discovery control in the previous experiment, also detected these six phyla. However, applying the two-sample *t*-test to the unnormalized data led to the detection of three different phyla. This discovery is consistent with previous results indicating that *Bacteroidota, Firmicutes, Actinobacteriota*, and *Fusobacteriota* are associated with obesity and that the obesity rate is significantly higher for immigrants than for people living in Thailand (Ley et al., 2006; Turnbaugh et al., 2009; Andoh et al., 2016). Furthermore, *Desulfobacterota* is shown to be related to inflammatory bowel diseases (Loubinoux et al., 2002), and the incidence of these diseases is much higher in Western countries compared to Asian countries, especially Thailand (Riansuwan and Limsrivilai, 2021), which is also consistent with our findings. These results again suggest that RSim normalization can improve the ability of differential abundance tests to control false discoveries and detect significant differences more effectively than existing normalization methods.

## 3 Discussion

In this study, we present RSim, a novel normalization method that corrects sample-specific bias in microbiome data with many zeros. RSim normalization is robust to the prevalent zeros because each step can work with zeros without making extra assumptions or treatments. RSim first identifies a set of non-differential abundant taxa by evaluating the pairwise rank similarity of taxa, then uses the estimated set as a reference set in reference-based normalization. This approach effectively corrects the compositional bias, even when microbiome data consists of many zero counts. Furthermore, while our discussion primarily focused on microbiome sequencing data, the ideas in this algorithm could potentially be applied to single-cell RNA sequencing data, where one major obstacle in normalization is also the zero counts problem.

Our comprehensive investigation into how normalization results can affect downstream analysis shows that the unobserved sampling fraction is a common confounder in high throughput sequencing data analysis. Compositional bias may confound the results of almost all types of downstream analysis, ranging from data visualization to statistical testing. This confounding factor creates false clusters or discoveries and obscure signals of interest in data analysis and interpretation. Our numerical experiments demonstrate that RSim normalization can eliminate compositional bias better than existing methods, reducing false discovery and increasing detection power in downstream analysis, including PCoA plotting, association analysis, and differential abundance analysis. We hope this new normalization method can improve the current data analysis pipeline and enable biological researchers to make more scientific discoveries.

One major assumption of RSim normalization is that more than half of the taxa are non-differential abundant, which is also used in developing differential abundance analysis in compositional data. This assumption may appear strong, but it is necessary for model identification when the sampling fraction is not observed (Wang, 2023b). In other words, the set of non-differential abundant taxa cannot be determined from the observed sequencing count when less than half are non-differential abundant. We recommend applying RSim normalization on high-resolution data, such as at ASV or OTU level, to satisfy this assumption. When ASVs/OTUs are aggregated into taxa at a higher taxonomic level, like class or order level, there could be much fewer non-differential abundant taxa because aggregation of non-differential and differential abundant ASVs results in differential abundant taxa (Wang, 2023a).

Finally, the development of RSim normalization shows that reference-based normalization can successfully correct compositional bias when identifying a set of non-differential abundant taxa. While RSim normalization only suggests one way to detect a set of non-differential abundant taxa, there could be alternative approaches to achieve the same goal. For example, Spearman ‘s rank correlation coefficient can be replaced by other correlation coefficients, such as the Pearson and Kendall rank correlation coefficients. It would also be interesting to explore if there is a better way to control the misclassification rate than our empirical Bayes method.

## 4 Methods

### 4.1 Reference-Based Normalization

In order to correct for compositional bias in sequencing data, various methods have been proposed in the literature for experiments both with and without spike-in bacteria. One of the most commonly used approaches is to calibrate the count data using the count of a control sequence, in cases where spike-in bacteria are available (Stämmler et al., 2016; Tourlousse et al., 2017; Tkacz et al., 2018; Hardwick et al., 2018). In experiments with spike-in bacteria, exogenous taxa of known concentration are introduced to each sample in equal amounts, and the count data is then rescaled using the count of these exogenous taxa. Specifically, suppose we have *n* samples and each sample has *d* taxa. Let *A*_*i,j*_ be the true absolute abundance of taxon *j* of the *i*th sample, and let *N*_*i,j*_ be the corresponding observed sequence counts. If we denote the collection of spike-in taxa as 𝒥*, then we can rescale the count data as follows:

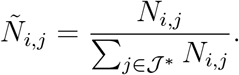

The aim of this scaling is to convert relative abundance to absolute abundance by ensuring that the rescaled count of the spike-in taxa is the same across samples. Experiments have shown that this scaling can successfully recover absolute abundance, with the error in recovery reduced by using multiple spike-in taxa (i.e., a larger 𝒥*) (Stämmler et al., 2016). However, using spike-in bacteria can be limited by the availability of reliable taxa, as well as potential amplification biases (Suzuki and Giovannoni, 1996; Brankatschk et al., 2012). Given these challenges, it is natural to ask whether this idea can be generalized to experiments without spike-in bacteria.

A new computational normalization method can be inferred from the way of scaling in experiments with spike-in taxa: first, we identify a data-driven reference set 𝒥_0_ whose absolute abundance remains stable across samples, and then normalize the count data with respect to this set:

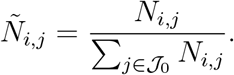

In the experiment with spike-in bacteria, the reference set is simply the set of spike-in taxa with the same absolute abundance across different samples. This normalization method is referred to as reference-based normalization in this paper. A reference-based approach is also widely employed in compositional data analysis (Aitchison, 1982; Pawlowsky-Glahn and Buccianti, 2011). For instance, the additive log-ratio transformation uses the last taxon as the reference set, while the centered log-ratio transformation uses the geometric mean of all taxa as the reference set. The reference-based hypothesis is also employed in differential abundance analysis of compositional data (Brill et al., 2022; Wang, 2023b,a). Unlike standard compositional data analysis, we utilize the sum of abundance in a set as the reference.

To perform efficient normalization in the absence of prior knowledge of spike-in taxa, we need to select an appropriate reference set, denoted by 𝒥_0_. In the presence of spike-in taxa, the reference set is simply the set of taxa with known absolute abundance that is constant across samples. However, for experiments without spike-in taxa, we can instead use a large set of non-differentially abundant taxa as the reference set. We assume that there exists a set of non-differentially abundant taxa called 𝒥_0_, such that their absolute abundance is similar across samples. When this is the case, the sum of the absolute abundance of these taxa is also similar across samples

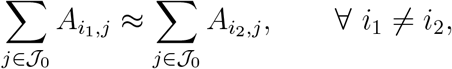

making the sum of abundance a suitable reference for normalization. Moreover, the sum of abundance of many taxa is generally more stable than that of a single taxon, due to the concentration of measure phenomenon (Talagrand, 1996; Boucheron et al., 2013). This observation suggests that normalization based on the set of non-differential abundant taxa can be effective in recovering the absolute abundance. In the next section, we will discuss how we estimate the reference set 𝒥_0_ from the data.

### 4.2 Reference Set Identification by Rank Similarity

In the previous section, we proposed using reference-based normalization to convert relative abundance to absolute abundance by identifying a large set of non-differential abundant taxa. In this section, we introduce a new method for detecting this set by comparing the count similarity between pairs of taxa. Before we present the method, we introduce some notation and assumptions. We partition the taxa into two groups based on absolute abundance: the collection of differential abundant taxa, denoted by 𝒥_1_, and the collection of non-differential abundant taxa, denoted by 𝒥_0_. To simplify the analysis, we assume that the absolute abundance of non-differential abundant taxa is the same across samples

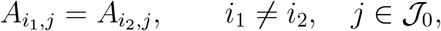

while the absolute abundance of differential abundant taxa varies between samples

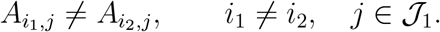

This model is only for illustrative purposes, but the method we introduce here can work in a more general setting, as long as the variance of absolute abundance for non-differential abundant taxa is much smaller than that for differential abundant taxa. In practice, we observe the count of each taxon, which only reflects relative abundance, and assume that it is drawn from a multinomial distribution

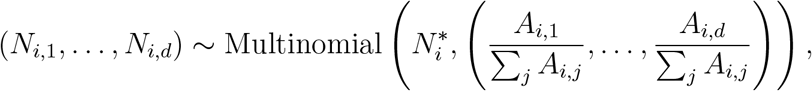

where 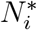 is the total sequence number in the *i*th sample. A similar model is also considered in Brill et al. (2022); Wang (2023a). Equivalently, we assume that the observed count of taxa is approximately equal to the absolute abundance multiplied by some unobserved sampling fraction *c*_*i*_

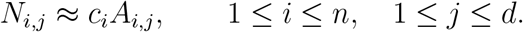

To make the model identifiable, we assume that |𝒥_0_| > *d/*2, where *d* is the number of taxa. See more discussion on model identification in Wang (2023b). After introducing these notations and assumptions, we present a two-step method for identifying the reference set.

#### Step 1: Differential Abundance Level Statistics

In the first step, we use the pairwise similarity of taxa to construct the statistics for the differential abundance level of each taxon (i.e., belonging to 𝒥_0_ or 𝒥_1_). The key observation we use in this step is that the observed count of two non-differential abundant taxa is much more similar than that of a non-differential abundant taxon and a differential abundant taxon. We represent the count of the *j*th taxon in all samples as 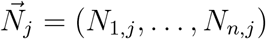 and the sampling fraction of all samples as 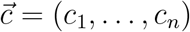. Since the absolute abundance of non-differential abundant taxa is stable across samples, we can expect that the count vectors of two non-differential abundant taxa, 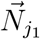 and 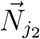, are almost proportional to the sampling fraction vector 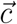, so the correlation between 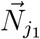 and 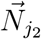 is close to 1. However, since the absolute abundance of differential abundant taxa varies across samples, we can expect the correlation between 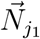and 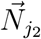to be much smaller than 1 when *j*_1_ is a non-differential abundant taxon and *j*_2_ is a differential abundant taxon. If we use Spearman ‘s rank correlation coefficient to measure correlation, then

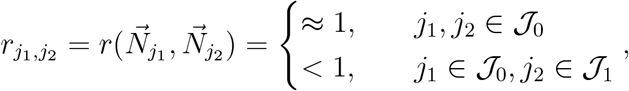

where *r*(*·, ·*) is Spearman ‘s rank correlation coefficient. How do we use the difference in the pairwise similarity to distinguish non-differential and differential abundant taxon? Since we assume that more than half of the taxa are non-differentially abundant, we can look at the median of the rank correlation coefficients between a taxon and other taxa. More concretely, if we denote the median as

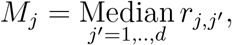

we can expect

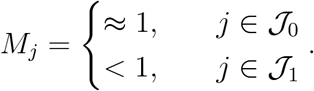

This observation suggests that the median *M*_*j*_ can be used to distinguish non-differential and differential abundant taxa.

#### Step 2: Taxa Classification

The second step of our method uses *M*_*j*_ to classify each taxon based on the empirical Bayes framework. The first step suggests that *M*_*j*_ is larger in non-differential abundant taxa than in differential abundant taxa. A natural classification rule is that we can choose a threshold *T* such that all taxa with *M*_*j*_ > *T* are classified as non-differential abundant taxa, i.e., 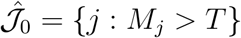 is the estimator for 𝒥_0_. The threshold *T* should help ensure that most taxa in 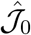 are non-differential abundant, since our goal is to find a reference set 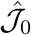 that satisfies the condition

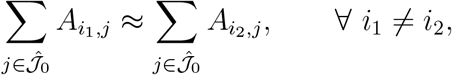

To achieve this, we choose the threshold *T* to control the misclassification error rate in 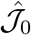, i.e.,

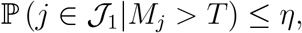

where *η* > 0 is the target misclassification rate that users select (e.g., *η* = 0.01). We estimate *T* using the empirical Bayes framework (Robbins, 1951; Efron, 2012). To facilitate this, we write *F*_0_ and *F*_1_ as the cumulative distribution functions of *M*_*j*_ when *j* is the non-differential and differential abundant taxa, respectively, and *F* as the cumulative distribution functions of *M*_*j*_, i.e.,

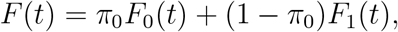

where *π*_0_ is the proportion of non-differential abundant taxa. Following these notations, we can rewrite the misclassification error in 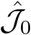 as

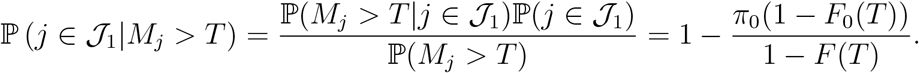

The idea in the empirical Bayes framework is to estimate *π*_0_, *F*_0_, and *F* from observed *M*_*j*_, *j* = 1, …, *d*, and then we can estimate the misclassification error by plugging in these estimators. The cumulative distribution function *F* can be naturally estimated by its empirical version

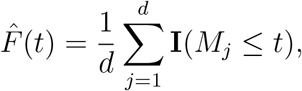

where **I**(*·*) is an indicator function. Before estimating *F*_0_ and *π*_0_, we choose *γ* > 0 such that ℙ(*M*_*j*_ > *γ*|*j* ∈ 𝒥_1_) ≈ 0, indicating that *j* is likely a non-differentially abundant taxon when *M*_*j*_ > *γ*. After choosing *γ*, we adopt a resampling method to estimate *F*_0_: 1) find all taxa with *M*_*j*_ > *γ* and define 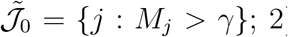 repeat subsampling the taxa from 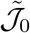 and recalculate the median on the subsampled data as in Step 1; 3) the empirical cumulative distribution function of these resampled medians is our estimator 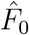. After finding 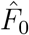 and 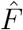, we can estimate *π*_0_ by

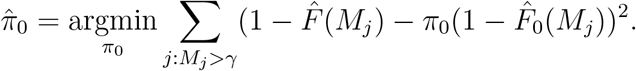

Here, we use the fact that 1 − *F*(*t*) ≈ *π*_0_(1 − *F*_0_(*t*)) when *t* > *γ*. With the estimators 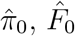, and 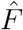 in hand, we choose the threshold 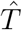 and estimated reference set as

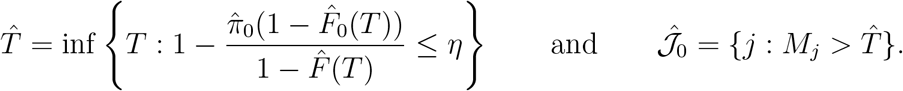

#### Choices of Tuning Parameters

RSim normalization has two main parameters: the target misclassification rate *η* and the threshold for differential abundance level statistics *γ*. The choice of *η* affects the empirical misclassification rate and the size of the estimated reference set, which in turn affects the performance of downstream analysis and sampling fraction recovery. A smaller *η* leads to a less significant bias but higher variance in sampling fraction recovery. Therefore, we recommend a smaller *η* for downstream analysis that is sensitive to sampling fraction recovery bias, such as differential abundance analysis. Conversely, a larger *η* is suitable for downstream analysis that requires an estimated sampling fraction with inflated bias and low variation, such as PCoA plotting.

The threshold *γ* should ideally be at the lowest level where statistics of differential abundant taxa cannot achieve. The choice of *γ* depends on the microbiome data characteristics, such as taxonomic rank and proportion of differential abundant taxa. Our experience suggests that at an ASV or OTU level, the statistics of at least 90% of non-differential abundant taxa are greater than 0.8. Therefore, we recommend using *γ* = 0.8 and use it in all experiments. As discussed, it is better to apply RSim normalization at an ASV/OTU level and then convert the data into higher taxonomic ranks, such as genus and family. Finally, note that RSim normalization performance is more sensitive to *η* than *γ*.

## Acknowledgments

The authors acknowledge support from NSF Grant DMS-2113458.

## Data Availability

All three data sets can be downloaded from Qiita (https://qiita.ucsd.edu/). Data set in He et al. (2018) is under study ID 11757. Data set in Vangay et al. (2018) is under study ID 12080. Data set in Caporaso et al. (2011) is under study ID 550.

## Code Availability

The R package is available at https://github.com/BoYuan07/RSimNorm. All analyses can be found under https://github.com/BoYuan07/RSim-manuscript-code.

## Supplement Material

### Numerical Experiment Setup

#### Summary of notations

We denote sample size by *n*, and the number of taxa by *d*. 𝒥_1_ represents the set of differential abundant taxa, and 𝒥_0_ represents the rest of the taxa. *X*_*i,j*_ is the raw count from the data set. *A*_*i,j*_ is the simulated absolute abundance. *N*_*i,j*_ is the simulated observed count. The parameter of misclassification rate control level is denoted by *η*. ℐ_1_ and ℐ_2_ represent two different groups of samples defined by the differential abundant taxa.

### Simulation study to assess compositional bias correction

The simulations in this section are conducted on the data set collected in He et al. (2018). We only include samples under the age of 30, so there are 539 samples and 37532 ASVs.

#### Setting for Figure 2

In subfigure (a), (b), and (c), *n* = 500 samples are randomly selected from the data set and randomly divided into two groups ℐ_1_ and ℐ_2_ with equal size. 10% taxa are randomly selected to be 𝒥_1_. The idea of our simulation experiments is to treat the raw count data as the population of absolute abundance. Specifically, absolute abundance is generated in the following way:

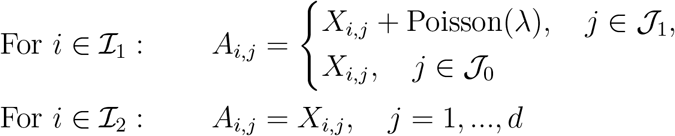

Given the simulated absolute abundance, observed count in the simulation experiment is generated in the following way:

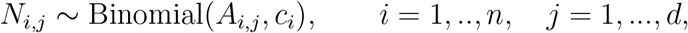

where *c*_*i*_ ∼ Unif[0, 1] is the sampling fraction of each sample. *λ* is 1 for the weak signal (a), 10 for the moderate signal (b), and 500 for the strong signal (c).

The setting for (d) is the same as (a), (b) and (c), except that we consider the proportion of differential abundant taxa as *p* = 0.1, 0.2, 0.3. The bias is evaluated in the following way:

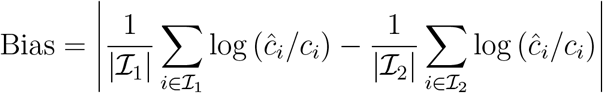

Here | *·* | represents the cardinality of a set. The two terms in above equation represent the average log difference between true and estimated sampling fraction for two groups respectively. If sampling fractions are correctly estimated for both two groups, the absolute difference between these two terms should be close to zero. On the other hand, if the differential abundant taxa lead to a systematic bias in estimated sampling fraction, the bias can be large. In all above experiments, we choose *η* = 0.2.

### Simulation study to assess the misclassification rate control

Similar to the last set of simulation experiments, we still conduct the experiments on data set collected in He et al. (2018). Each set of experiment is repeated 500 times and we use the average misclassification rate as measure.

#### Setting for Figure S1a

*n* = 500 samples are randomly selected from the data set and divided into two groups ℐ_1_ and ℐ_2_ with equal size. 10% taxa are randomly selected to be 𝒥_1_. Absolute abundance is generated in the following way:

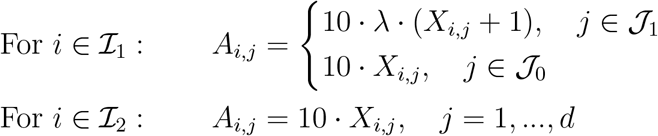

*λ* is the signal strength and set to be 2 in this experiment. Observed count is generated in the following way:

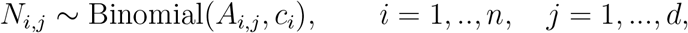

where *c*_*i*_ ∼ Unif[0, 1] is the sampling fraction of each sample.

#### Setting for Figure S1b

*n* = 500 samples are randomly selected from the data set and divided into two groups ℐ_1_ and ℐ_2_ with equal size. The top 10% abundant taxa are selected to be 𝒥_1_. Absolute abundance is generated in the following way:

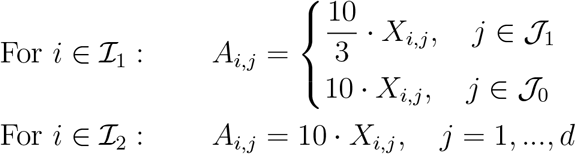

Observed count is generated in the following way:

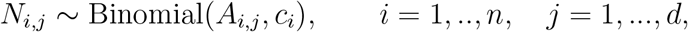

where *c*_*i*_ ∼ Unif[0, 1] is the sampling fraction of each sample.

#### Setting for Figure S1c

*n* = 500 samples are randomly selected from the data set. The top 10% abundant taxa are selected to be 𝒥_1_. First we generate a random variable *Y* = [*Y*_1_, .., *Y*_*n*_] and *Y*_*i*_ ‘s are drawn from Unif[1, 100] independently. And absolute abundance is generated in the following way:

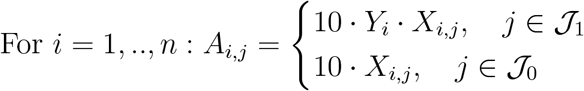

Observed count is generated in the following way:

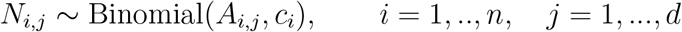

where *c*_*i*_ ∼ Unif[0, 1] is the sampling fraction of each sample.

#### Setting for Figure S2

All the four subfigures share the same data structure with Figure S1a. Signal strength *λ* increases from 1 to 10; the ratio between the group sizes increases from 0.1 to 0.4; the proportion of differential abundant taxa increases from 0.1 to 0.5; sample size increases from 100 to 500. We choose *η* = 0.01 in the experiments.

### Simulation study for PCoA

The Euclidean distance is used in all the PCoA plots.

#### Setting for Figure 3a and 3b

The experiment is conducted on data set collected in He et al. (2018). We only include samples with total counts larger than 30000 in this experiment. Figure 3a is the result for raw data. For 3b, half of the samples are subsampled to 1/10 of the original sequencing depth, while the other half of the samples remain the same. *η* = 0.2 in these experiments.

#### Setting for Figure 3c

PCoA performance on the data set collected in Vangay et al. (2018). The setup of this experiment can be found in Section 2.3.

#### Setting for Figure S3a

All the samples of the data set in He et al. (2018) are included in this experiment. Samples are divided into two groups ℐ_1_ and ℐ_2_. The top 25% abundant taxa are selected to be 𝒥_1_. Absolute abundance is generated in the following way:

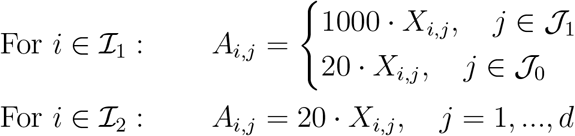

Observed count is generated in the following way:

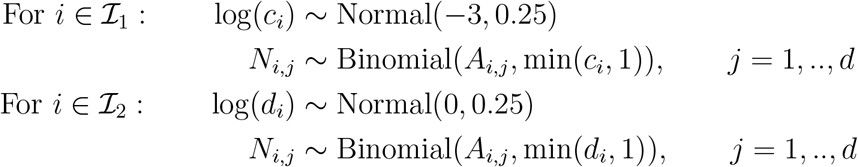

The sampling fraction of ℐ_1_ is around ten times of ℐ_2_.

#### Setting for Figure S3b

All the samples are included in this experiment(*n* = 539). The top 1% abundant taxa are selected to be 𝒥_1_. First we generate a random variable *Y* = [*Y*_1_, .., *Y*_*n*_] and *Y*_*i*_ ‘s are drawn from Unif[5, 500] independently. Absolute abundance is generated in the following way:

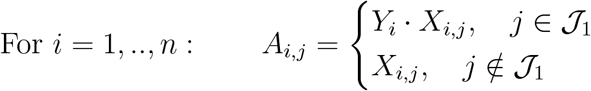

Observed abundance is generated in the following way:

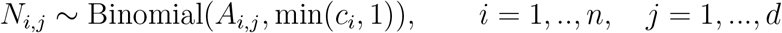

where 1*/c*_*i*_ ∼ *N*(*Y*_*i*_, 1) and min(*c*_*i*_, 1) is the sampling fraction for sample *i*.

### Simulation study for association analysis

Simulation experiments is still based on the data set in He et al. (2018). All the experiments for the association analysis are repeated 500 times.

#### Setting for Figure 4a

*n* = 500 samples are divided into two groups with equal size. Samples in the first group are subsampled to 1*/c* of the original sequencing depth, while the second group keeps the same as the original data set. *c* increases from 1 to 3. We choose *η* = 0.07.

#### Setting for Figure 4b

We consider sample size *n* = 100, 300, 500. Samples are divided into two groups ℐ_1_ and ℐ_2_ with equal size. The top 10% abundant taxa are selected to be 𝒥_1_. Absolute abundance is generated in the following way:

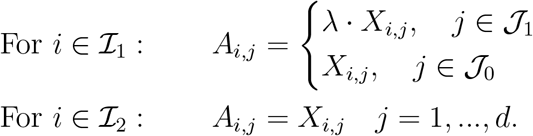

Here we choose *λ* = 2. Observed count is generated in the following way:

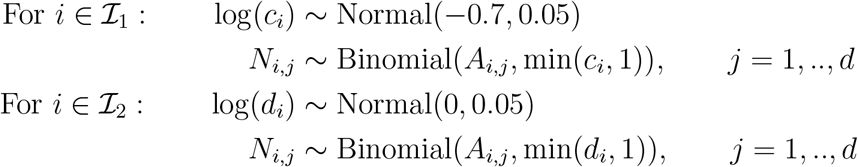

The sampling fraction of ℐ_1_ is around two times of ℐ_2_. We choose *η* = 0.05.

#### Setting for Figure

The setting of Figure S5a is the same as Figure 4b except that *λ* = 5.

#### Setting for Figure S5b

We consider sample size *n* = 100, 300, 500. The setting is the same with Figure S1c.

### Simulation study for differential abundance test

There are three parts in this section. The first set of experiments is based on the data set in He et al. (2018), of which results are shown in Figure 5; the second set is based on the data set in Caporaso et al. (2011), of which result is shown on Figure 6; the third set is based on the data set in Vangay et al. (2018), of which result is shown on Table S1. We use Benjamini-Hochberg procedure to adjust the effect of multiple testing.

For the first set of the simulation study, to reduce the computational load, we select the subtree below Node 351 as our target, which contains 1081 taxa in total. We repeat each experiment 500 times and consider the average sensitivity rates and false discovery rates.

#### Setting for Figure 5a and 5b

We consider sample size *n* = 100, 200, 300. Samples are divided into two groups ℐ_1_ and ℐ_2_ with equal size. 10% taxa are randomly selected to be 𝒥_1_. Absolute abundance is generated in the following way:

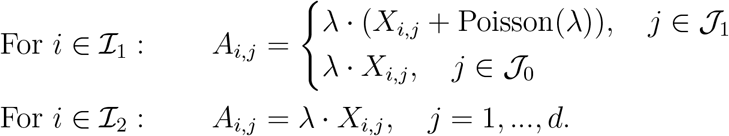

Observed count is generated in the following way:

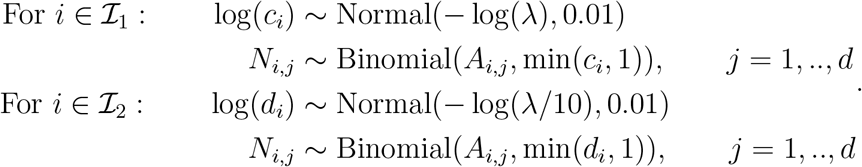

*λ* is chosen to be 50,100,150. Two-sample *t*-test is applied on the normalized counts. The significance level is 0.1. We choose *η* = 0.01.

#### Setting for Figure 5c and 5d

We consider sample size *m* = 200, 350, 500. The top 10% abundant taxa are selected as 𝒥_1_. First we generate a random vector *Y* = [*Y*_1_, .., *Y*_*n*_] and *Y*_*i*_ are draw from Unif[1, *λ*]. The absolute abundance is generated in the following way:

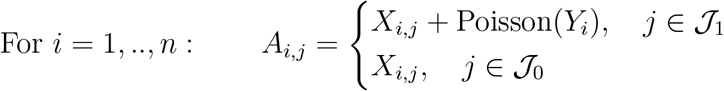

Then we generate the sampling fraction in the following way:

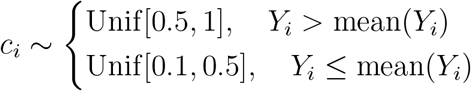

Observed abundance is generated in the following way:

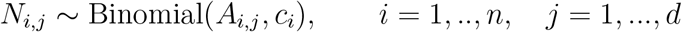

*λ* is chosen to be 50, 100, and 150. Pearson correlation test is applied on normalized counts. The significance level is 0.1 and we choose *η* = 0.01.

#### Setting for Figure 6

Samples of subject M3 ‘s right palm are selected from data set in Caporaso et al. (2011). In total there are 352 samples and 24333 ASVs. Samples with a sequencing depth of less than 10000 are selected to be the first group, and samples with a sequencing depth of more than 20000 are another group. ANCOM, EdgeR, LinDA, RDB and *t*-test with different normalization methods are considered in the experiment. We use the detected 0.9 for ANCOM, which is the most conservative setting, and the default setting for EdgeR, RDB and LinDA.

#### Setting for Table S1

Differential abundant test is conducted for group KareThai and group Karen1st from data set in Vangay et al. (2018). For *t*-test with RSim, data is first normalized at the ASV level, then aggregated to the phylum level. The FDR control level is 0.1.

## Extra Numerical Results and Figures

**Figure S1:**
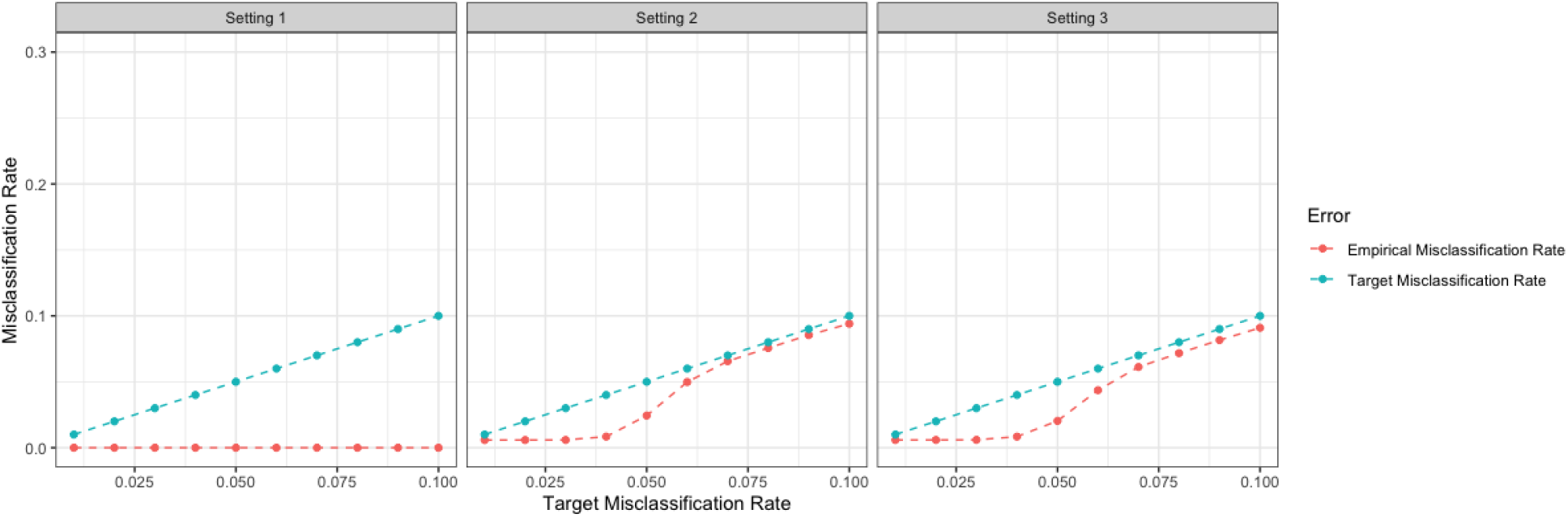
Misclassification rate control when the target misclassification rate *η* varies. In all figures, the *x*-axis is the target misclassification rate while the *y*-axis represents the empirical misclassification rate of the estimated reference set. Setting 1: 10% taxa are randomly selected as differential abundant taxa, and the latent variable of differential abundant taxa is binary; Setting 2: the differential abundant taxa are top 10% most abundant taxa, and the latent variable of differential abundant taxa is binary; Setting 3: the differential abundant taxa are top 10% most abundant taxa and the latent variable of differential abundant taxa is continuous. In all settings, the misclassification error rate of the estimated reference set can be well controlled.

**Figure S2:**
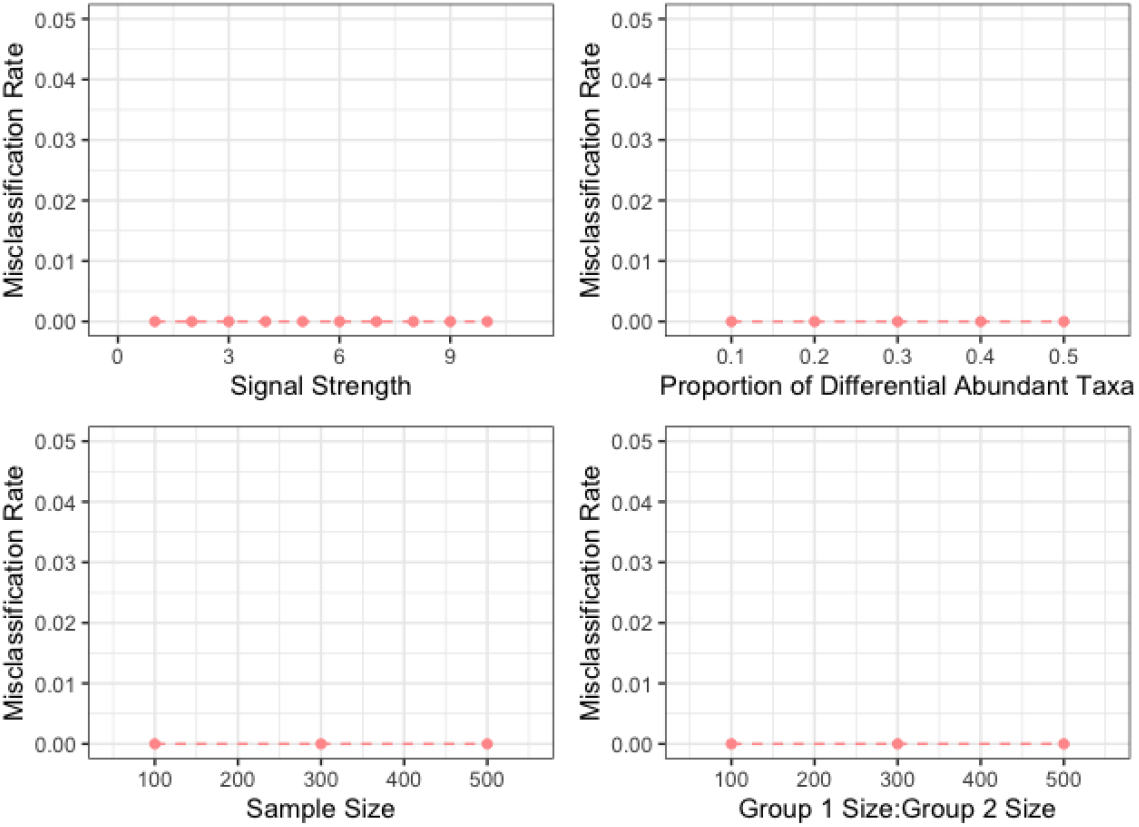
Misclassification rate control when the signal strength of differential abundant taxa, the balance of group size, proportion of differential abundant taxa, and sample size are different. The empirical misclassification rate in RSim is well controlled despite the choices of the signal strength of differential abundant taxa, the balance of group size, proportion of differential abundant taxa, and sample size.

**Figure S3:**
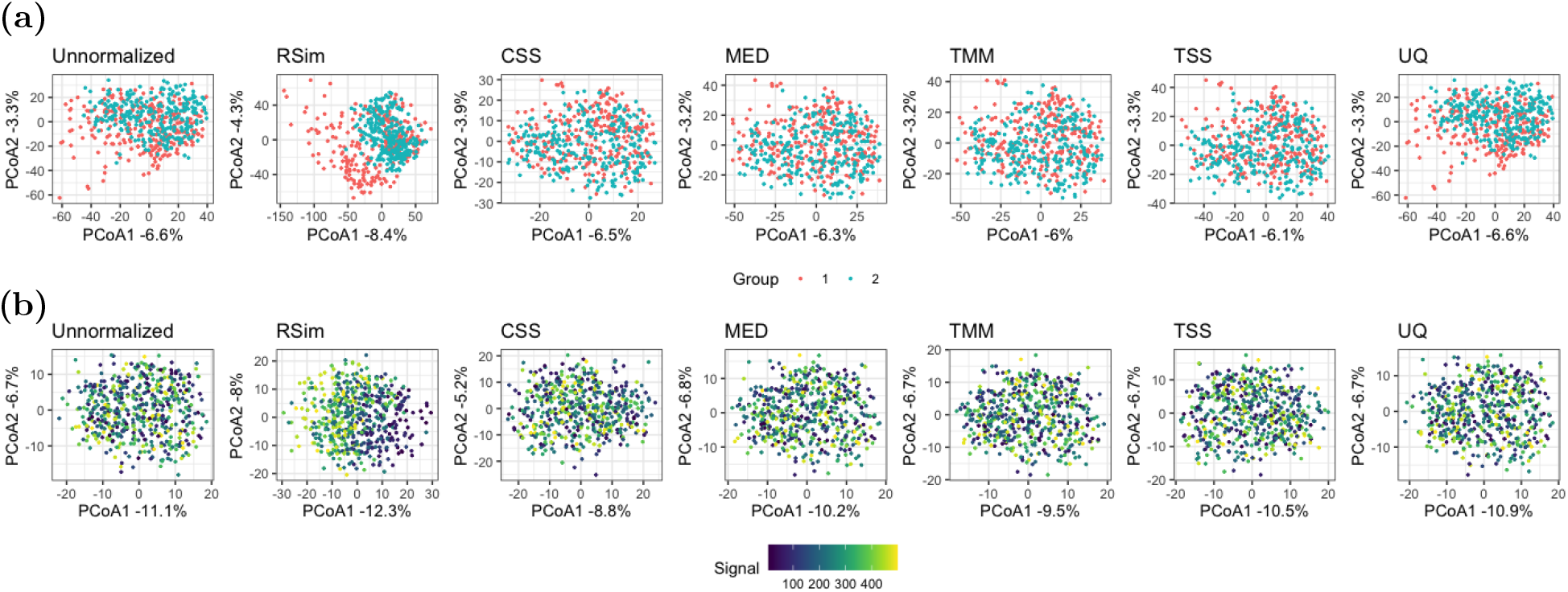
Normalization can reveal biological pattern in PCoA plots. In (a), samples are randomly divided into two groups, and the top 10% most abundant taxa are differential abundant taxa with a binary latent variable. In (b), the top 10% most abundant taxa are differential abundant taxa with a continuous latent variable. In these figures, RSim normalization can reveal the structure of the latent variable. Euclidean distance with log transformation is used in all PCoA plots.

**Table S1:** Differential abundant phyla detected by different differential abundance analysis methods. Three methods are considered: *t*-test on unnormalized data, *t*-test on data normalized by RSim, and RDB test on unnormalized data.

| Method | Phylum |
| --- | --- |
| *t*-test on unnormalized data | Cyanobacteria |
|  | Elusimicrobiota |
|  | Proteobacteria |
| *t*-test on data normalized by RSim | Actinobacteriota |
|  | Bacteroidota |
|  | Desulfobacterota |
|  | Firmicutes |
|  | Fusobacteriota |
|  | Patescibacteria |
| RDB test on unnormalized data | Actinobacteriota |
|  | Bacteroidota |
|  | Desulfobacterota |
|  | Firmicutes |
|  | Fusobacteriota |
|  | Patescibacteria |

**Figure S4:**
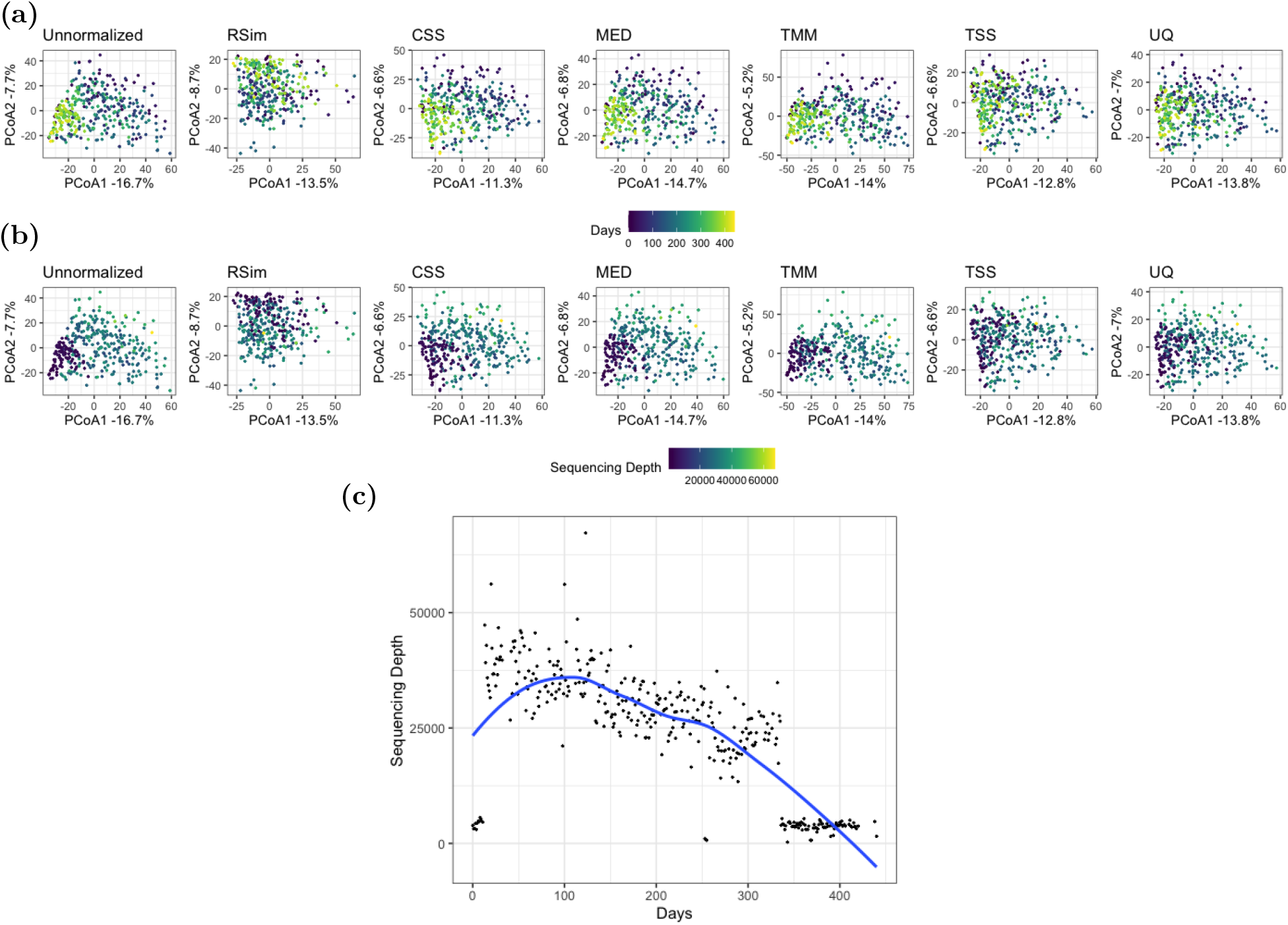
False pattern caused by compositional bias leads to a misleading conclusion. (a) shows the PCoA plots colored by days after the experiment started. (b) presents the PCoA plots colored by sequencing depth. (c) show the relationship between time and sequencing depth. The pattern of time in PCoA plots is highly overlapped with pattern of the sequencing depth, which can be explained by the deterministic relationship between time and sequencing depth.

**Figure S5:**
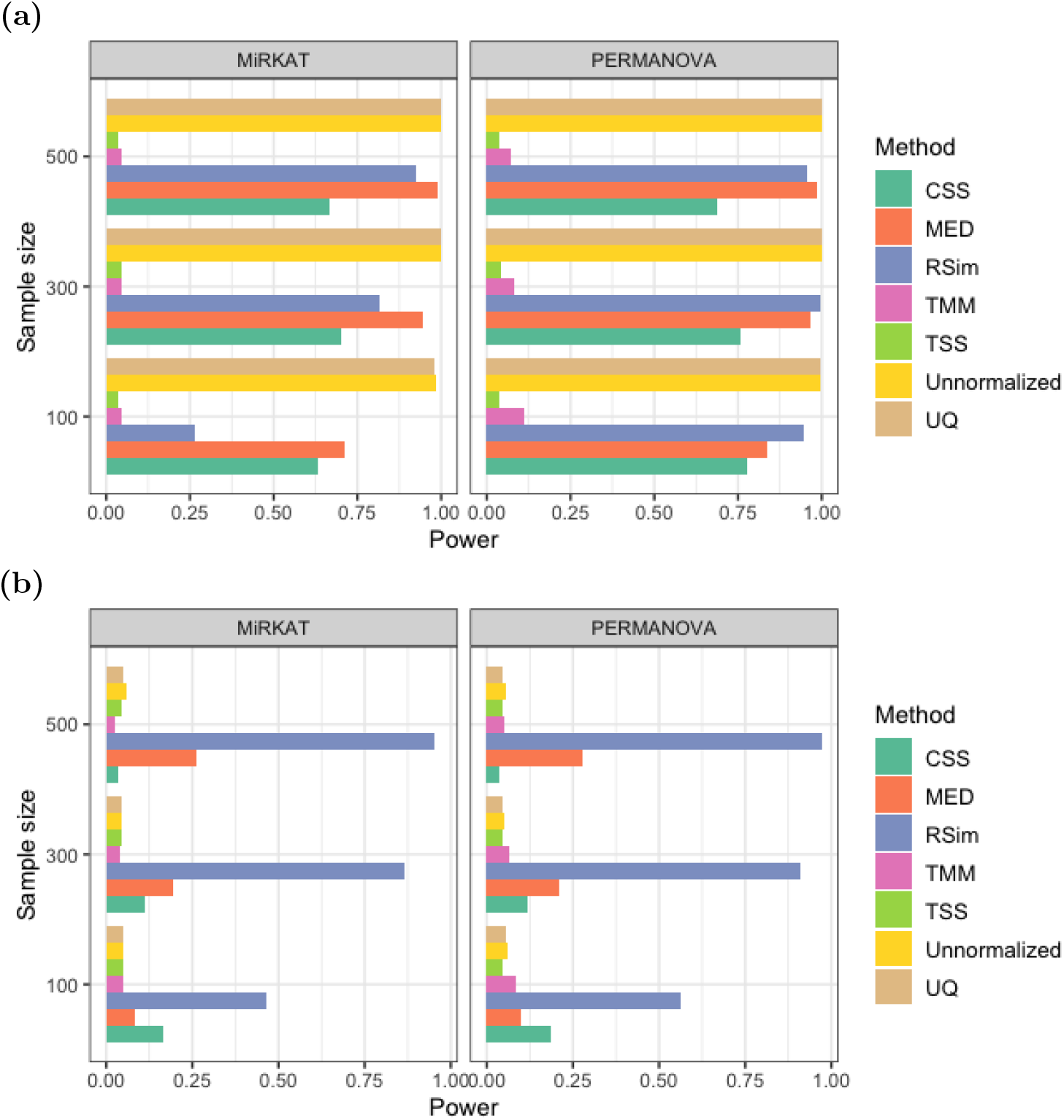
Normalization can improve the power of association analysis. In (a) and (b), samples are randomly divided into two groups, and the top 25% most abundant taxa are differential abundant taxa with a binary or continuous latent variable. The significance level is 0.05. RSim can improve the power of association analysis.

